# Multiple lineages, same molecular basis: task specialization is commonly regulated across all eusocial bee groups

**DOI:** 10.1101/2020.04.01.020461

**Authors:** Natalia de Souza Araujo, Maria Cristina Arias

## Abstract

A striking feature of advanced insect societies is the existence of workers that forgo reproduction. Two broad types of workers exist in eusocial bees: nurses which care for their young siblings and the queen, and foragers who guard the nest and forage for food. Comparisons between this two worker subcastes have been performed in honeybees, but data from other bees are scarce. To understand whether similar molecular mechanisms are involved in nurse-forager differences across distinct species, we compared gene expression and DNA methylation profiles between nurses and foragers of the buff-tailed bumblebee *Bombus terrestris* and of the stingless bee *Tetragonisca angustula*. These datasets were then discussed comparatively to previous findings on honeybees. Our analyses revealed that although the expression pattern of genes is often species-specific, many of the biological processes and molecular pathways involved are common. Moreover, DNA methylation and gene expression correlation were dependent on the nucleotide context.

## Introduction

Caste specialization in eusocial insects is a notorious example of polyphenism, where multiple morphological and behavioural phenotypes emerge from the same genotype^1,2^. In social Hymenoptera (bees, wasps and ants), queen and worker reproductive castes perform distinct functions in the colony. While queens undertake reproductive duties, workers perform all the other tasks necessary for nest maintenance and growth^3^. Two broad categories of workers exist in eusocial bees: nurses and foragers^4,5^. Nurses are responsible for comb construction, offspring/queen care and internal colony maintenance, while foragers perform tasks related to external colony defence and resources provisioning^5,6^. In advanced eusocial bee species, such as honeybees, worker subcastes are mainly age determined; younger bees are nurses and when they become older, they switch to being foragers^7,8^. In primitively eusocial species, such as the social bumblebees, specialization in worker subcastes is not so straightforward^9,10^.

To investigate differences in bee worker subcastes, many studies have been conducted in the highly eusocial honeybee (*Apis*). Gene expression comparisons have identified expression changes between worker behaviours^1,5,7,11,12^, which could even be used to predict neurogenomic states in individual bees^13^. Similarly, profiles of DNA methylation, an epigenetic marks that likely underpins gene expression differences, were additionally shown to directly correlate with worker task^14,15^. Certain genes were differentially methylated according to the worker subcaste and foragers that were forced to revert to nursing restored more than half of the nursing-specific DNA methylation marks^16,17^.

Many of the molecular differences between honeybee workers and nurses could have arisen later in the evolution of this lineage. To broadly understand how subcastes evolved it is necessary to differentiate such more recent changes – that could be species-specific – from those shared across species, and thus likely ancestral. The highly eusocial stingless bees have age-based division of labour^18^, similarly to that of honeybees despite their most common ancestor being 50 to 80 million years ago^19,20^. To date, no global expression or epigenetic studies have been performed in stingless bees to understand worker task specialization. Similarly, while primitively eusocial bumblebees are largely studied ecological biological models and important wild and managed pollinators, we know comparatively little about the molecular underpinnings of differences between its worker subcastes. Indeed, studies have been restricted to few genes, leaving many open questions^21–23^. A major limiting element for these studies is that this species display a somewhat fluctuating division of labour with indistinctive separation between subcastes^10,21,23^.

We aim to fill in this knowledge gap through the analyses of the global gene expression differences between nurses and foragers, and the characterization of nurses DNA methylation profile in two eusocial bee species, the primitively eusocial buff-tailed bumblebee, *Bombus terrestris*, and the highly eusocial stingless bee, *Tetragonisca angustula*. Combined, these two bee species and the honeybee represent the three evolutionary branches of eusocial corbiculates sharing a common social origin^24^. Hence, in addition to using the generated datasets to uncover unique and more recent molecular traits linked to task division in *B. terrestris* and *T. angustula*, we also verified whether common genes and pathways could be involved in task specialization across all the eusocial bee groups.

## Results

### Reference transcriptome assemblies

For both species we used as reference a transcriptome set of superTranscripts^25^, in which multiple transcripts from the same gene are represented in a single sequence. *B. terrestris* workers had 27,987 superTranscripts of which 431 are potentially lncRNAs and 21,638 (77,3%) were annotated. The final *T. angustula* assembly had 33,065 superTranscripts, and was largely complete. Indeed, 26,623 superTranscripts (80.5%) had a high sequence similarity to known protein-coding genes from other species in the UniRef90 database, and 347 were considered lncRNAs (transcriptomes available at https://github.com/nat2bee/Foragers_vs_Nurses). A summary of major quality parameters from the two species datasets can be found on Table SI.

### Differential expression analyses in <u>Bombus terrestris</u>

Since task division in *B. terrestris* workers is a plastic behaviour^9,21^, we performed a principal component analysis of the normalized read counts as an additional verification step to validate our sampling method. The main components clearly clustered nurses and foragers samples separately (Figure S1) indicating that our sampling method was efficient to obtain two distinct groups in bumblebee workers, here considered as nurses and foragers due to the activities they were performing when sampled. We found 1,203 differentially expressed superTranscripts between the two worker groups (Figure S2), whereby 436 superTranscripts were more highly expressed in nurses (Supplementary file S2) and 767 were more highly expressed in foragers (Supplementary file S3). The majority of these superTranscripts (77.3% and 72.6% respectively) have similarity to known protein-coding genes, while respectively three and one are possible long non-coding RNAs (lncRNAs). Five Gene Ontology (GO) biological processes terms (“transposition”, “DNA-mediated, transposition”; “DNA integration”; “DNA recombination”; and “pseudouridine synthesis”) were overrepresented among the differentially expressed superTranscripts (p < 0.01; Table SII).

### Differential expression analyses in <u>Tetragonisca angustula</u>

In workers of *T. angustula* 241 superTranscripts were differentially expressed between nurses and foragers (Figure S2). Among these, 179 had higher levels of expression in nurses, being 157 genes with a significant blast hit to protein databases (Supplementary file S4). Foragers had 62 superTranscripts reported as more highly expressed than in nurses of which 59 were annotated (Supplementary file S5). Enrichment analyses revealed 30 GO terms for biological process (BP) as enriched in the tested set of differentially expressed superTranscripts when compared to the entire transcriptome (p < 0.01; Table SII), including processes related to mitochondrial metabolism (“aerobic respiration”; “respiratory electron transport chain”; “oxidative phosphorylation” and “mitochondrial ATP synthesis coupled electron transport”) and other metabolic process (“lipid metabolic process” and “carbohydrate metabolic process”).

### DNA methylation in worker genes

Whole bisulfite sequencing (WBS) from *B. terrestris* and *T. angustula* nurses were used to screen DNA methylation patterns in the entire transcriptome and among the differentially expressed superTranscripts. Because *T. angustula* lacks a reference genome and most DNA methylation reported in bees occur within gene exons^14^, we performed methylation analyses by mapping bisulfite sequenced reads to the transcriptomes and not genomes (complete estimations available at https://github.com/nat2bee/Foragers_vs_Nurses). In *B. terrestris* 23.14 % of all cytosine sites are in CG (cytosine/guanine) context. This is a higher proportion than in *T. angustula* where 15.44 % of all C sites available occur in CG context. We find this explains the higher proportion of CG methylation observed in the bumblebee (Figure 1). Nevertheless, in both species DNA methylation in CG context was enriched, that is there was more DNA methylation in CG context than it would be expected simply based on the proportion of sites available. Furthermore, superTranscripts general methylation (mC) levels are higher in *T. angustula* (mean mC 1.24 %) than in *B. terrestris* (mean mC 0.66 %) (Figure 2).

**Figure 1.**
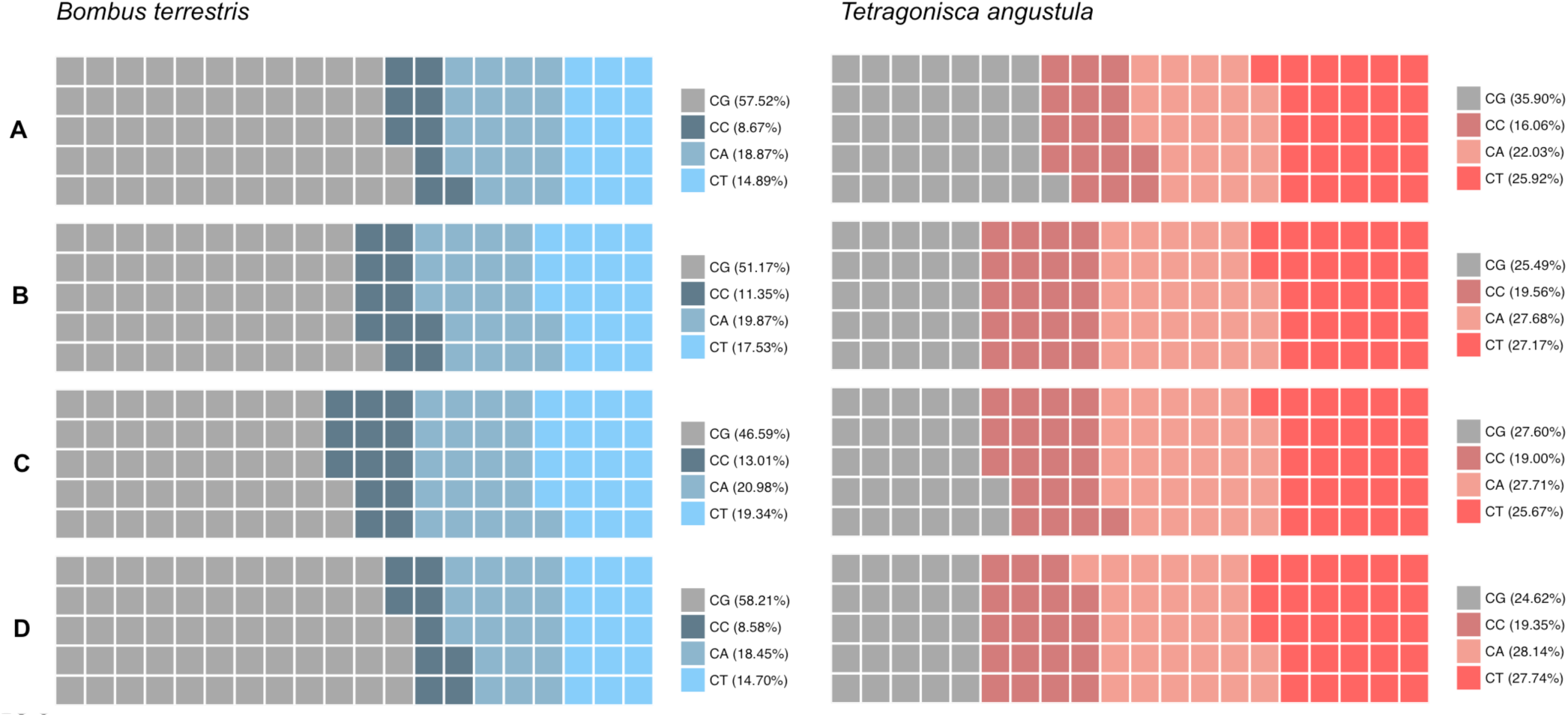
Nucleotide context in which the methylated cytosines occur proportionally to all methylated cytosines reported in nurses of *B. terrestris* and *T. angustula*, in distinct gene sets. **A** – in the entire transcriptome; **B** –in the differentially expressed superTranscripts between foragers and nurses; **C** – in the superTranscripts with higher expression levels in foragers; **D** – in the superTranscripts with higher expression levels in nurses. Grey squares represent methylation at CG context; methylation in non-CG context is illustrated in different shades of blue for *B. terrestris* and in red shades for *T. angustula*. One square ≈ 1%, considering all mC reported sums up to 100%.

**Figure 2.**
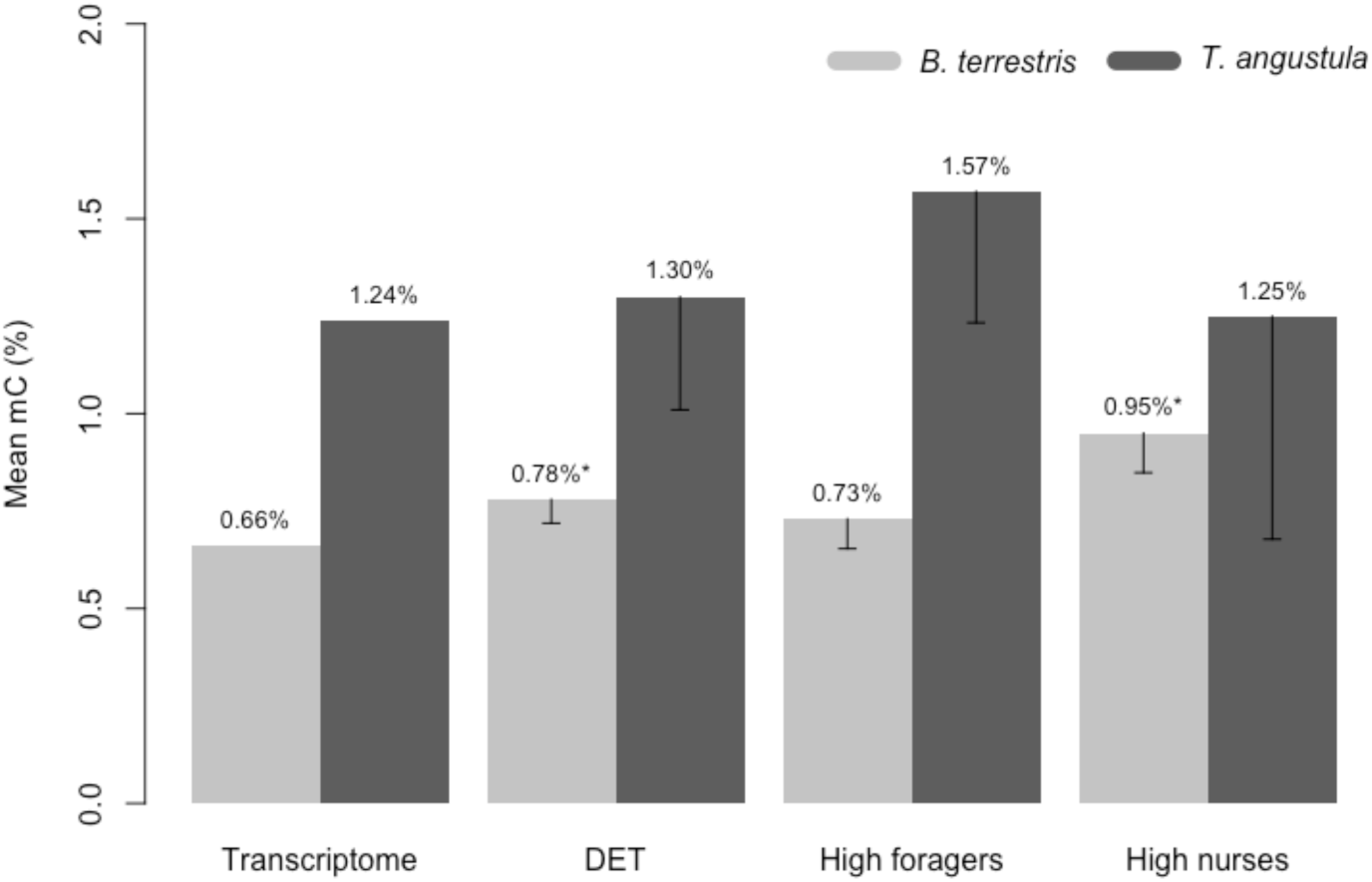
Mean mC levels in distinct gene sets of *B. terrestris* and *T. angustula* nurses. Transcriptome – refers to the values observed in the complete transcriptome; DET – differentially expressed superTranscripts between nurses and foragers; High foragers – superTranscripts with higher expression levels in foragers when compared to nurses; High nurses – superTranscripts with higher expression levels in nurses when compared to foragers. ^**\***^ significantly different from the global transcriptomic mean, with p < 0.01 at 95% CI; confidence interval bars are shown.

In both species the differentially expressed superTranscripts had higher levels of methylation than the overall transcriptomic mean (Figure 2), however only in *B. terrestris* this difference was significant (*B. terrestris* p = 6.267e-4, *T. angustula* p = 0.3669 at 95% CI). Interestingly, while in *B. terrestris* this increase was mostly due to the greater methylation level of superTranscripts highly expressed in nurses; the mean mC level of the highly expressed superTranscripts in *B. terrestris* nurses was 43.93% higher than the global transcriptomic mean (p = 1.339e-06 at 95% CI). In *T. angustula* superTranscripts highly expressed in foragers were the more methylated ones (Figure 2), although still not at a significant level when compared to the general mean (p = 0.05355 at 95% CI). The nucleotide context in which the methylated cytosines occurred also varied in each gene subset (Figure 1). There was an overall reduction in the contribution of CG methylation in the subset of differentially expressed superTranscripts when compared to the entire transcriptome, except for superTranscripts highly expressed in *B. terrestris* nurses (Figure 1D).

Combined these findings suggest a correlation between mC and gene expression depending on the methylation context. Indeed, we identified a positive correlation between global transcript expression levels and CG methylation at both species (*B. terrestris* r_s_ *=* 0.23 and *T. angustula* r_s_ = 0.24) but not with CW (CA – cytosine/adenine or CT – cytosine/thymine) methylation (*B. terrestris* r_s_ *=* 0.08 and *T. angustula* r_s_ = −0.07). Curiously, when we used only the set of differentially expressed superTranscripts, no correlation was found between gene expression and mC in *B. terrestris*, neither in CG (r_s_ = 0.08) nor in CW (r_s_ = −0.06) context. However, in *T. angustula*, both types of methylation correlated negatively with gene expression in this scenario (CG r_s_ = −0.31; CW r_s_ = −0.35). This suggests that DNA methylation indeed plays a role in subcaste task division of other eusocial bee species, as in honeybees, but in a more complex way than previously recognized.

### Comparative analyses of genes involved in task division in the two species

In order to recognize species-specific from shared molecular mechanisms, different strategies were used. First, we asked whether the exact same genes were commonly involved in the observed subcaste differences of *T. angustula* and *B. terrestris*. Comparing the two sets of differentially expressed superTranscripts we identified 15 genes in common (Table I; Figure 3C), which is significantly more than it would be expected by chance (p = 6e-04, mean number of genes expected 7.04, SD=2.58). Interestingly, the expression pattern of these genes was not always equivalent in both species (Table I). Seven genes were commonly highly expressed in nurses of both species when compared to foragers, (p = 4e-04, mean number of genes expected 2.16, SD=1.45), but only two genes were commonly highly expressed in foragers (p = 0.3062, mean number of genes expected 1.1, SD=1.03).

**Figure 3.**
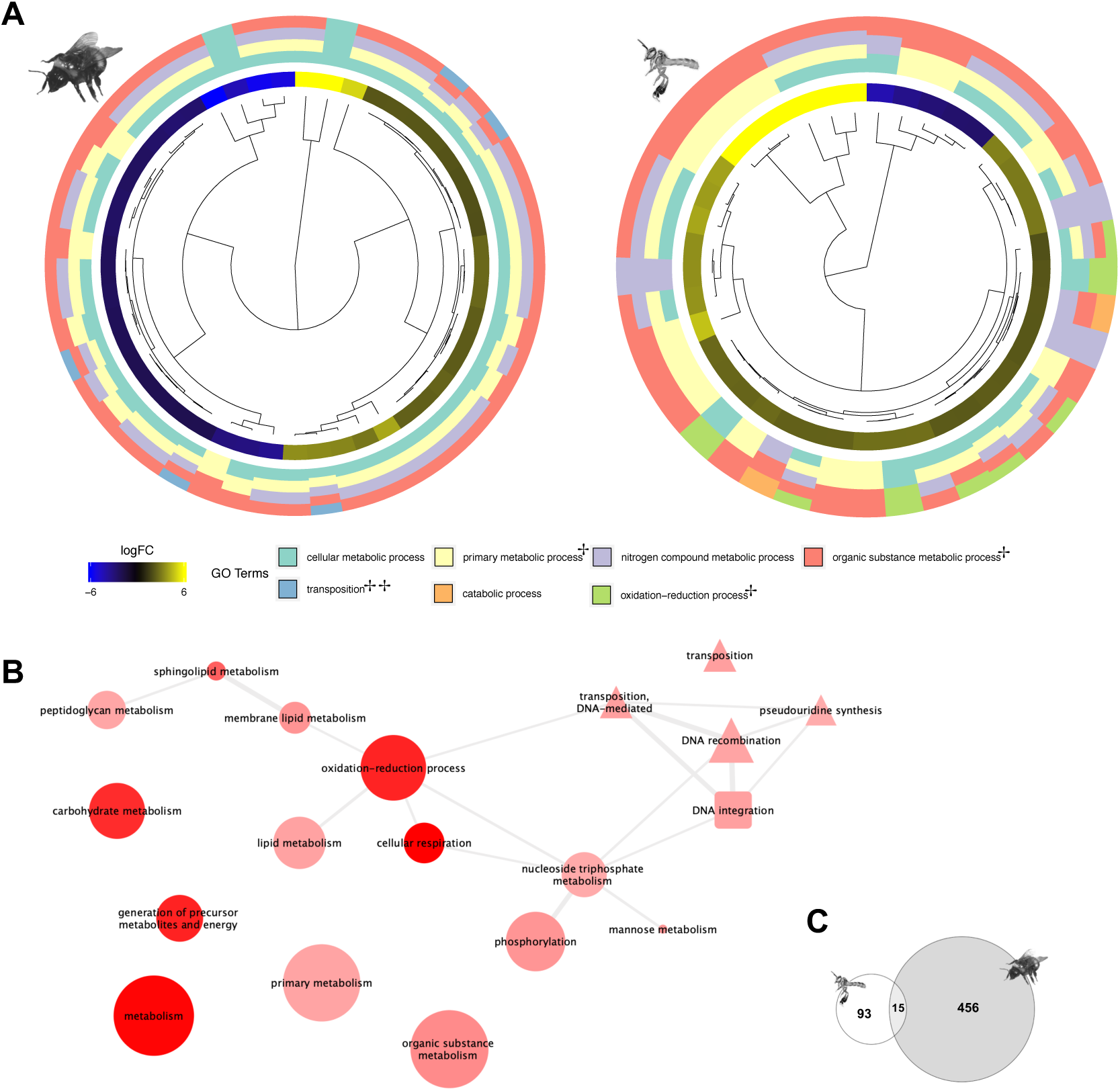
Comparisons between *B. terrestris* and *T. angustula* GO processes involved in task specialization. **A –** Hierarchical clustering of the differentially expressed superTranscripts with the third hierarchical level of GO annotation organized by their mean logFC difference between nurses and foragers. Outer circle colours show to which GO term the gene could be associated to. ^‡‡^ BP term enriched in the set of differentially expressed superTranscripts of *B. terrestris*; ^‡^ BP term enriched in the set of differentially expressed superTranscripts of *T. angustula*. **B** – Similarity network of the enriched GO terms, after semantic similarity-based reduction. GO terms that are very similar to each other are linked and the line width indicates the degree of similarity. Edge shape indicates whether the shown term is enriched in *B. terrestris* (circle), in *T. angustula* (triangle) or in both species (square). Edge colour intensity indicates the p-value in the enrichment test (the darker the colour tone, the smaller the p-value). Edge size indicate the frequency of the GO term in the entire UniProt database. **C** – Euler diagram showing the number of genes in common between the set of differentially expressed superTranscripts of each species.

**Table I.**
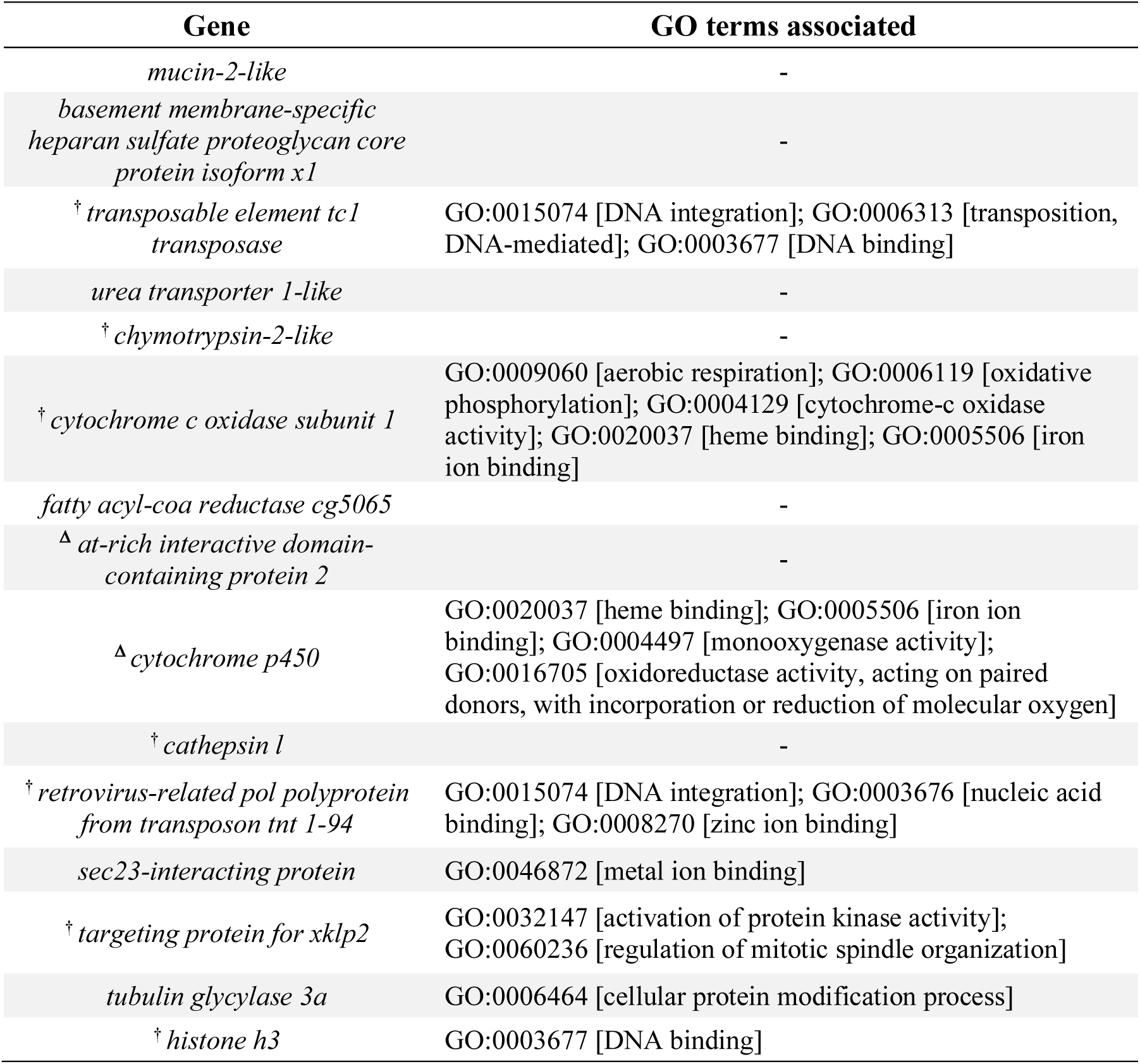
List of genes in common to the sets of differentially expressed superTranscripts between nurses and foragers of *T. diversipes* and *B. terrestris*. ^**†**^ Indicate genes commonly highly expressed in nurses of the two species when compared to foragers. ^**Δ**^ Indicate genes commonly highly expressed in foragers of the two species when compared to nurses.

Secondly, we investigated whether the same molecular pathways could be involved in the task division of the two species. For this, we searched for similarities among the biological processes to which the differentially expressed superTranscripts were related. We used a comparative approach based on GO subgraphs of the enriched terms. This type of subgraph relies on the hierarchical graphic structure among GO terms, where parent terms are more general and less specialized than child terms^26,27^. Consequently, using subgraphs it is possible to compare not only the enriched terms themselves but also their hierarchical connections, reducing gene annotation bias^28^. In this comparison (Figures S3, S4 and S5) we found that the enriched GO terms of the two species were associated and eventually all of them nested under two main processes (Figure S3): “metabolic process” (GO:0008152) and “cellular process” (GO:0009987). Thus, although specific enriched terms are distinct in both species (only “DNA integration” is commonly enriched), this divergence disappears at the parental levels of the topology and almost all terms in *B. terrestris* subgraph are also contained in *T. angustula* subgraph (Figure S3). At the third hierarchical level (Figure 3A), lineage specific GO processes start to emerge such as “transposition” (GO:0032196) in *B. terrestris*, and “catabolic process” (GO:0009056) and “oxidation-reduction process” (GO:0055114) in *T. angustula*. Nevertheless, superTranscripts showing the greatest differences in expression within species (i.e. higher absolute mean logFC between groups) are not the ones related to these species-specific processes (Figure 3A). The connection between the enriched GO terms in the set of differentially expressed superTranscripts of *B. terrestris* and that of *T. angustula* can also be visualized on semantic similarity-based clusters (Figure 3B).

## Discussion

The comparison of present findings in *B. terrestris* and *T. angustula* with previously published information about task specialization in *Apis* workers are summarized on Table II. Because the literature about this topic in honeybees is extensive and these studies applied distinct methodologies of sampling, expression estimation and data analyses, we restricted our comparisons to a review of genes and molecular pathways commonly highlighted across studies.

**Table II.**
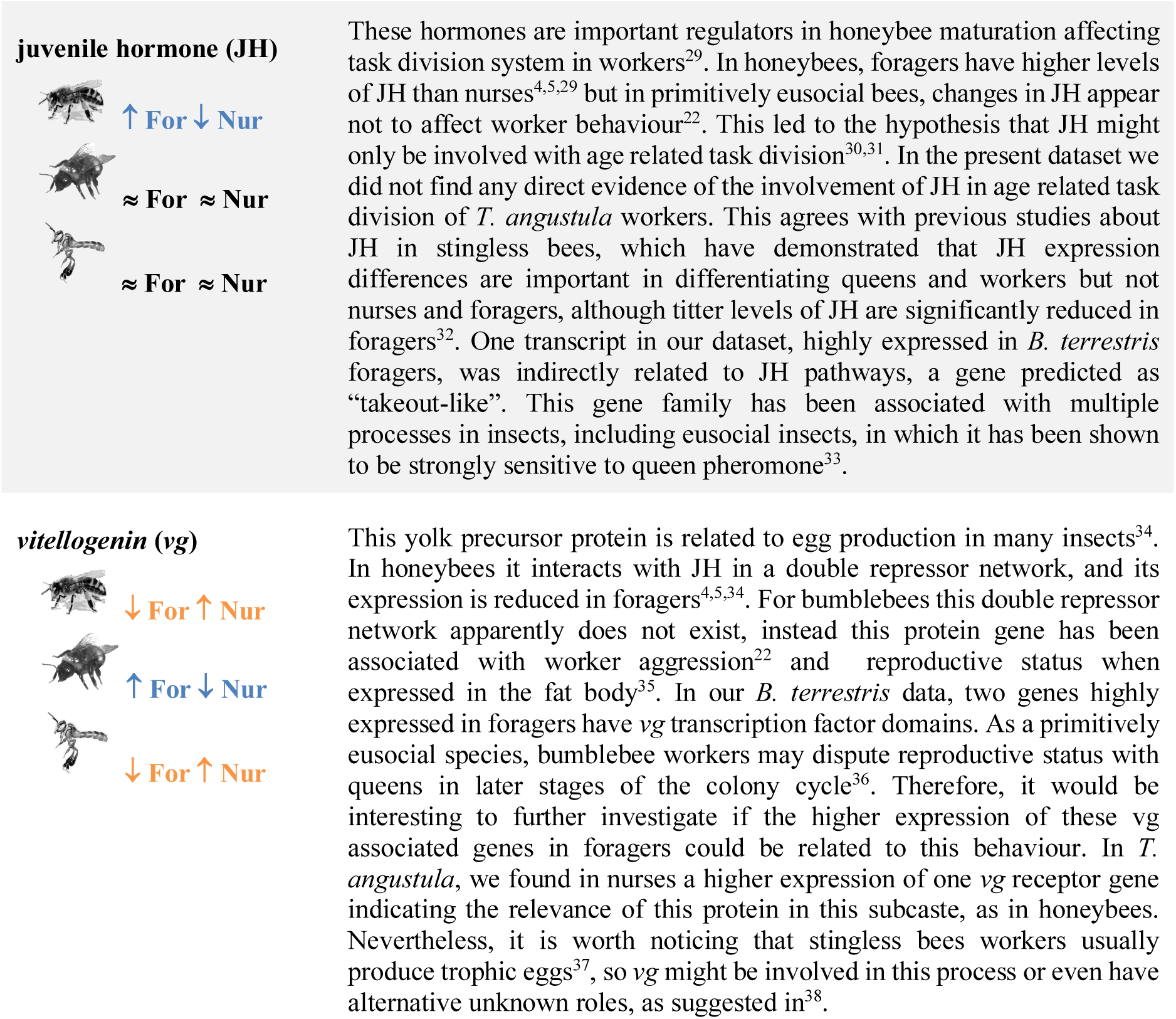
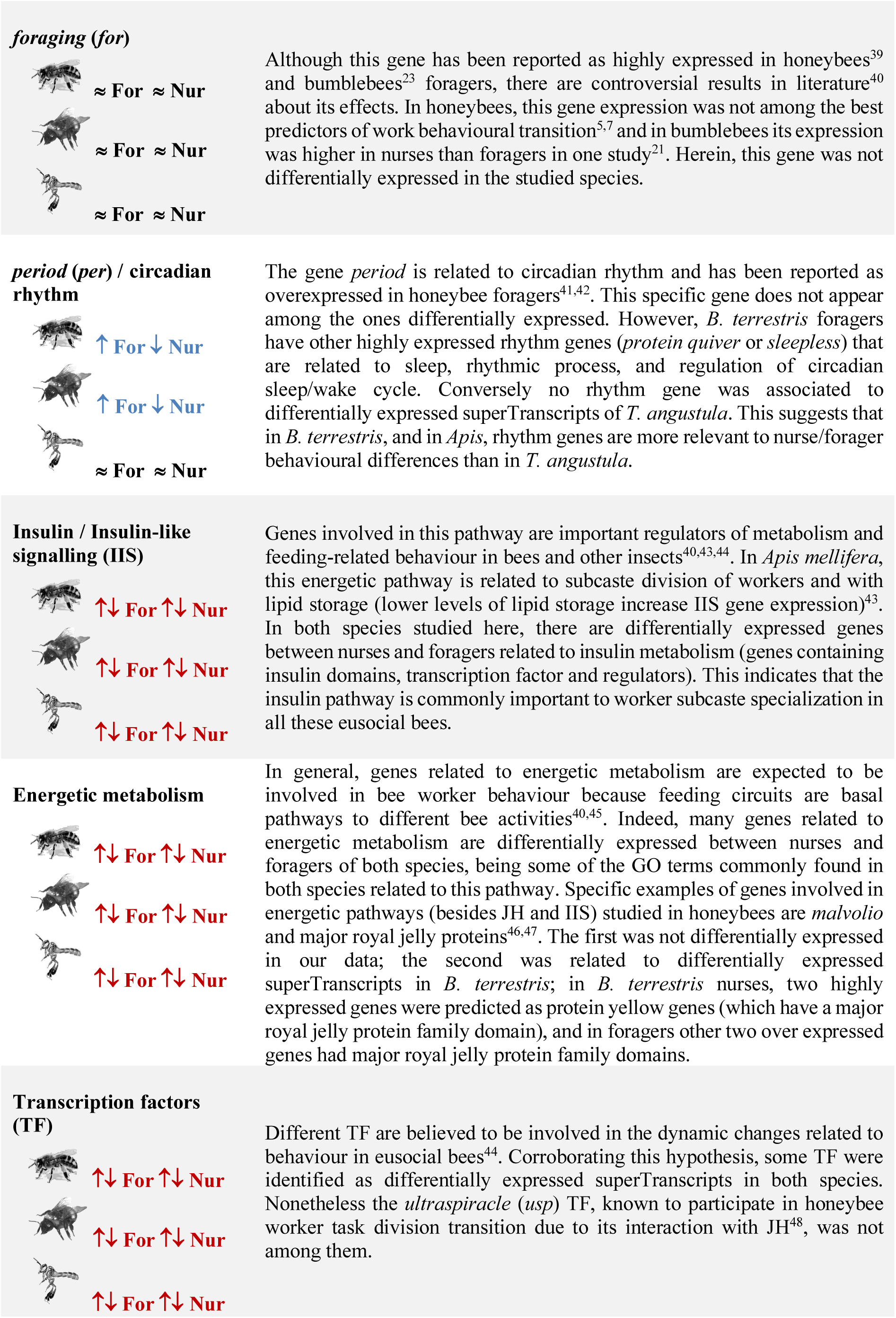
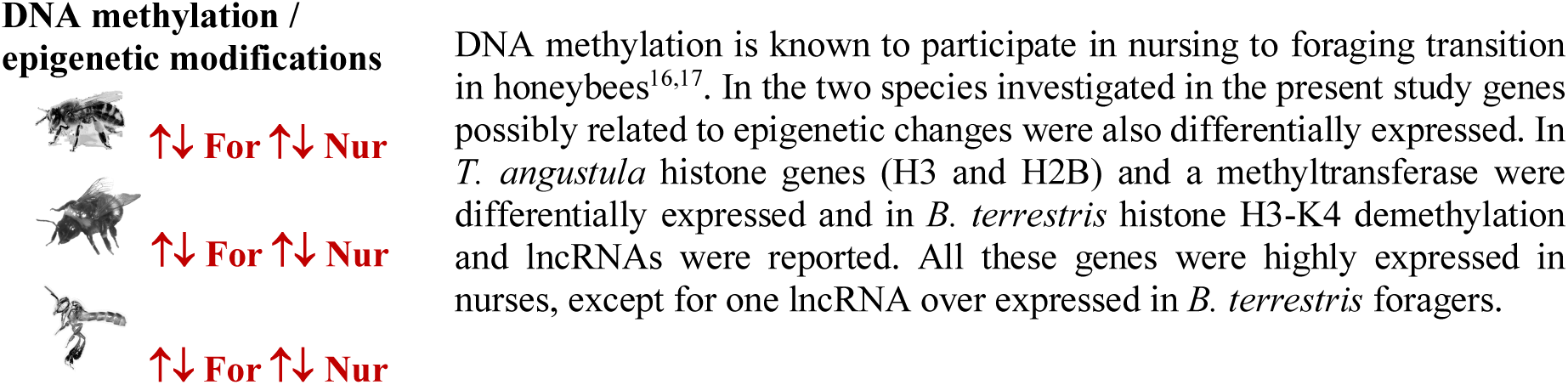
Genes and molecular pathways commonly highlighted in literature as involved in honeybee worker task division compared to present findings in *B. terrestris* and *T. angustula*. For – foragers; Nur – Nurses. Symbols indicate if: evidences suggest that the expression is higher (↑) or lower (↓) in one group compared to the other, in blue if foragers have higher expression levels than nurses and in orange if the opposite occurs; (≈) in black, no changes identified or controversial evidences; and (↑↓) in red, indicate a mixed pattern, with some genes in the pathway highly expressed and others with reduced expression level in one of the two subcastes.

Our comparisons sought to differentiate species-specific from common molecular mechanisms involved in worker task division across all eusocial lineages of corbiculate bees. Species-specific mechanisms were mostly related to the expression pattern of genes. Many of the differentially expressed genes were not common to all species, and among the ones that were, the pattern of expression was not necessarily the same. Genes highly expressed in one species subcaste were often down expressed in the same subcaste of the other species. For instance, genes related to circadian rhythm are highly expressed in foragers of *B. terrestris* and *Apis*^41,42^, but not in *T. angustula* foragers. Moreover, genes related to yolk production, such as *vg* related genes, are commonly highly expressed in nurses of *T. angustula* and *Apis* ^4,5,34^, but not in *B. terrestris* nurses. This is not unexpected since each lineage forgo unique selective pressures, despite presenting similar behaviours^49^. Even closely related species (within the same taxonomic genus) are known to differ in the expression pattern of certain genes^11^. This implies that the expression profile of particular genes, identified through the study of one single species, should not be directly extrapolated to explain other species responses.

A clear illustration of how misleading these assumptions can be is the *vg/*JH network, which has been largely studied in honeybees. Honeybee nurses present higher levels of *vg* and lower levels of JH when compared to foragers. Once workers become foragers their levels of JH increases, which in turn represses the *vg* expression in a double repressor network^4,48^. In bumblebees, as demonstrated previously^22^ and corroborated by our data, this network is not regulated in the same manner. In this bee, genes related to JH and *vg* are both highly expressed in foragers. For *T. angustula* we found supporting evidence of the involvement of *vg* in nursing behaviour, nonetheless JH genes were not highly expressed in foragers. This supports the hypothesis that in stingless bees the typical *vg/*JH double repressor network observed in honeybees is also not functional, and *vg* is distinctly regulated^32,38^.

Nevertheless, gene expression dynamics in worker behaviour is not completely unrelated across eusocial bees. Beyond the literal expression trend, we still found a significant number of genes commonly differentially expressed between nurses and foragers of *B. terrestris* and *T. angustula*. Interestingly, some of these common genes were also shown to be responsive to queen pherormone^33^. Genes like *cytochrome p450, fatty acyl-CoA*, mitochondrial and histone related were found to be sensitive to queen pheromone in ants and bees^33^. Moreover, biological processes terms enriched in each species set of differentially expressed superTranscripts were highly comparable. Our comparisons of enriched GO terms subgraphs highlighted broader similarities between *B. terrestris* and *T. angustula*, indicating how distinct GO terms (and genes) were involved in similar biological processes. In general, biological terms related to energetic and metabolic processes (“organic substance metabolic process”; “primary metabolic process”; “nitrogen compound metabolic process”; and “cellular metabolic process”) were central to subcaste differentiation of both species.

The relevance of metabolic pathways to insect sociality has been demonstrated in many studies over the years^45,50–52^ and it is most certainly not a species-specific trait. These pathways are affected by queen pheromone in different species and are involved with caste determination of multiple hymenopteran lineages, including bees, ants and wasps^33,53^. Given the central role of energetic and metabolic maintenance in any living animal it is not surprising that changes in these pathways will affect a number of features, including behavioural phenotypes. It is however fascinating to observe how plastic and dynamic, in terms of gene regulation, these networks can be, with different lineages frequently evolving unique responses but still being sensible to similar cues (like queen pheromone).

This mosaic pattern of species-specific features involved in common molecular processes is also observed in the epigenetic machinery. Transcriptomic and WBS data support the involvement of DNA methylation and other epigenetic factors in worker specialization of the two analysed species. Genes involved in epigenetic alterations were found among the differentially expressed superTranscripts of *T. angustula* and *B. terrestris*, and the species global methylation patterns were distinct from that of their differentially expressed superTranscripts. The differentially expressed superTranscripts had overall more mC (Figure 2) and less CG methylation (Figure 1). Still, a closer investigation revealed distinct specific epigenetic mechanisms in each species. To begin with, epigenetic related genes that are differentially expressed in each species are different. Likewise, only in *T. angustula*, the genes highly expressed in foragers were more methylated at CG context and had higher mean mC levels than genes overexpressed in nurses. The opposite was found in *B. terrestris*. Considering that WBS data was obtained from nurses in both species, these distinct patterns are quite unexpected.

Although in the studied species DNA methylation was frequent in CG context, methylation within other nucleotides contexts also occurred (i.e. non-CG or non-CpG methylation). Non-CG DNA methylation is frequently associated with a number of processes in plants^54,55^ and only recently its function in other eukaryotes have gained more attention^56^. Still, the effects of differential DNA methylation contexts in most organisms are poorly understood and underestimated (reviewed in^56,57^). Methylation in CG and non-CG sites are typically mediated by distinct mechanisms^58^; CG methylation is constitutively maintained by DNA methyltransferase 1 (Dnmt1)^56,57^, while non-CG methylation are kept by mechanisms of *de novo* methylation involving the DNA methyltransferase 3 (Dnmt3)^59^. Therefore, non-CG methylation is majorly related to new and more variable epigenetic alterations^57^. Supporting evidence for the existence of non-CG methylation in social insects was previously reported for ants^60^ and honeybees^59^. In honeybees, non-CG methylation seems to be involved with alternative mRNA splicing and is especially enriched in genes related to behavioural responses. However, no direct correlation with sociality could be stablished^59^. Herein, such correlation is demonstrated with the different proportions of CG and non-CG methylation observed in the set of differentially expressed superTranscripts when compared to the general transcriptomic profile. This indicate that both CG and non-CG methylation interplay in worker task division. Further data is needed to infer how specific methylation contexts could affect certain behavioural changes but based on the results gathered so far, we hypothesize that non-CG methylation dynamics is relevant to task division and possibly to other social traits.

Higher levels of mC in bees have been associated to an increase in gene expression, i.e. genes with more methylation also have higher expression levels^14^. We found this correlation to be true for CG methylation in both species tested, but not for methylation in non-CG context. In fact, among the differentially expressed superTranscripts of *T. angustula*, where greater levels of non-CG methylation are observed, we found a negative correlation between gene expression and DNA methylation. This suggests that the effect of mC in bee gene expression might also be dependent of the methylation context; CG methylation seems to increase gene expression while non-CG methylation might supress it.

Finally, it is important to consider some of the limitations of the present study. First, aiming to obtain a global overview of gene expression and DNA methylation differences we used full bodies for the transcriptomic and bisulfite sequencings. Since we know that different body parts, tissues and even cells have unique gene expression dynamics^12^ it is likely that our approach had reduced our power to detect small scale alterations and specific contexts. Moreover, to facilitate the comparisons between the two bees we used similar pipelines for them. This means that sometimes we compromised the bumblebee analysis to match it with the analysis of the species with no reference genome available. For example, we annotated both species transcriptomes based on search similarities to databases instead of using *B. terrestris* genome for its annotation. This approach might especially affect GO enrichment analysis. Differently from genome annotation, transcriptomic annotation is redundant, i.e. multiple transcripts (or superTranscripts in our case) may annotate to the same gene and this affects the frequency of GO terms in the dataset. To deal with this, we kept the frequency of GO terms proportional in the enrichment test by using the appropriate background list (in our case the complete transcriptome set), which is the used and recommended approach for GO enrichment tests^61^. However, our enrichment stats might still be biased by the chosen approach. Nevertheless, since GO annotations are dynamic and always biased by database representation^62^, we have chosen to keep the same methodological approach for both species. In this manner, if the enrichment test is biased it will be equally biased in both species facilitating comparisons. Finally, we did not validate our gene expression results with an alternative independent method (such as real time reverse polymerase chain reaction). Given due consideration, the present study can only describe broad patterns and conclusions regarding the species general expression and methylation profiles.

Further works should address detailed and more subtle differences. Through the analyses of the global transcriptomic and DNA methylation profiles of subcastes from two eusocial bee lineages, we gather an important dataset for the study of social behaviour evolution. These data aligned to a review of the honeybee literature, allowed comparisons among all eusocial corbiculate bee groups; Apini, Bombini and Meliponini. Main findings support the hypothesis that common and more ancient molecular mechanisms are involved in worker task division across these species, standing as central among them energetic and metabolic pathways, and epigenetic factors. However, despite these similarities, particular gene expression patterns tend to be species-specific. This scenario could be explained by later specialization of species-specific molecular responses to ancient social cues which left a mosaic profile in worker task division, where unique and shared features are found. Moreover, results indicate that non-CG methylation is relevant to worker behavioural dynamics and that it might affect gene expression differently from CG methylation. As a result, the involvement of non-CG methylation in other social traits should be further investigated.

## Material and Methods

### Sample collection and sequencing

Bee species were chosen based on their behaviour (primitively eusocial and highly eusocial), phylogenetic relationship (corbiculate bees^24^), and sampling convenience. Samples were from three colonies per species. *B. terrestris* colonies were obtained from commercial suppliers (Biobest®) and kept in lab condition at Queen Mary University of London (England). All bees in the colonies were marked and housed in wood boxes attached to foraging arenas. After 16 days of adaptation all recently born workers received an individual number tag; individuals used in the analyses were all tagged. Bumblebee workers usually do not forage right after emergency^63^, therefore we waited for five more days before start sampling. For *T. angustula*, colonies regularly kept at the Laboratório de Abelhas (University of São Paulo – Brazil) were used for sample collection.

Workers subcaste were determined in two different ways. First, for *B. terrestris*, nurses were selected based on observation. Colonies were observed for one day during all their active foraging period (6 hours uninterruptedly). Tagged bees who stayed inside of the nest during the entire period, never entering the foraging arenas, were considered nurses. In the following day, nurses and foragers were collected and immediately frozen in liquid nitrogen. Foragers were sampled first, while collecting nectar in the foraging arena. Nurses were posteriorly collected inside of the colonies. Then, for *T. angustula*, nurses were defined by age. Brood cells (close to emergency) were removed from the colonies and transferred to an incubator with controlled temperature and humidity. Upon emergency, female workers were marked with specific colours using a water-based ink and immediately returned to the colony. Ten to twelve days after their emergency and reintroduction, colonies were opened and marked individuals were sampled. During this age worker bees from *T. angustula* present nursing behaviour^37^. Foragers were collected while leaving and returning to the colonies from foraging trips. To prevent sampling of guard workers^2^, bees standing in front of the colony entrance were avoided. Some of the foragers were collected before nurse sampling and others after this period, but no foragers were sampled while nurses were marked and collected so as to avoid effects of colony disturbance in the worker behaviour. Nurses from different colonies were collected in different days.

All individuals were sampled between 10h-12h for both species, and entire worker bodies were used for RNA and DNA extraction. For RNA-Seq, six *T. angustula* workers, from the same colony and subcaste, were pooled as one sample. *B. terrestris* samples were a pool of RNA extractions from three workers per subcaste/ colony. Each colony was considered as one sample replicate. Total RNA was extracted from workers using Qiagen® extraction kit (RNeasy Mini Kits). RNA quality and quantification were verified using the Bionalyzer®, Nanodrop® and Qubit®. Samples were posteriorly used for RNA sequencing on Illumina® HiSeq 2000. Library preparation was performed by sequencing providers. *B. terrestris* workers were sequenced by the Genome Center at Queen Mary University of London, and *T. angustula* samples were sequenced at LACTAD (Unicamp). RNA sequencing generated 30-50 million paired reads (100bp) per colony replicate. For whole bisulfite sequencing, total DNA from one nurse (whole body) per species was used for the phenol-chloroform DNA extraction^64^. WBS were performed following the protocol described in^65^ using the Illumina® NextSeq500. WBS returned 60-70 million single reads (150 bp) per sample. Sequencing and library preparation were performed at University of Georgia. All sequenced reads are available at BioProject ID PRJNA615177.

### Transcriptome assembly and differential expression analyses and comparisons

Reads quality assessment was performed using the FastQC program^66^ (v0.11.2) before and after cleaning. The FASTX Toolkit^67^ (v0.0.14) was used to trim the first 14 bp of all reads because an initial GC bias^68^ was detected. Low quality bases (phred score below 30) and small reads (less than 31 bp) were removed using SeqyClean^69^ (v1.9.3). Samples from nurses and foragers were combined for the assemblies. To increase *de novo* transcriptome assembly efficiency, cleaned reads were digitally normalized^70^ (20x coverage). Transcriptome assembly were performed differently for each species. For *B. terrestris*, its genome^71^ was used as reference in two approaches. First, using HISAT2^72^ (v2-2.0.3) and StringTie^73^ (v1.2.2) a regular reference assembly was obtained. Secondly, the Trinity^74^ (v2.1.1) program was used to perform a reference guided *de novo* assembly. The two resulting assemblies were merged using CD-Hit^75^ (v4.6), Corset^76^ (v1.05) and Lace^25^ (v0.80) to cluster transcripts into superTranscripts. We have chosen to use this combined approach for *B. terrestris* for two reasons. First, to optimized the transcriptome assembly based on our dataset, a recommended procedure even for species with well-annotated reference genome and transcriptome^77^. Second, to make *B. terrestris* and *T. angustula* datasets more comparable since for the later we have used the clustering method. There is no reference genome for *T. angustula*, therefore we performed a combined *de novo* assembly using two strategies with the Trinity pipeline: a reference guided *de novo* assembly, based on the genome of another stingless bee, *Melipona quadrifasciata*^78^; and a complete *de novo* assembly. Afterwards, the two assemblies were merged as in the bumblebee. Assemblies used programs default recommended parameters, CD-Hit was used to merge transcripts with more than 95% similarity, Corset was set to keep transcripts with a minimum of 50x coverage, and Lace was used to obtain the superTranscripts.

SuperTranscripts were then annotated with Annocript^79^ (v1.2) using the UniProt Reference Clusters (UniRef90) and the UniProtKB/Swiss-Prot databases^80^ (June 2016 version). SuperTranscripts with significant blast hits (e-value < 1e-5) against possible contaminants (plants, fungus, mites and bacteria) in the UniRef90 were removed from the final datasets. Finally, only potentially coding superTranscripts (based on blast results and ORF analysis) or possible lncRNAs were kept. This annotation pipeline was used for both species. Quality parameters from the transcriptomes were analysed using QUAST^81^ (v4.0), BUSCO^82^ (v2), TransRate^77^ (v1.0.3) and Qualimap^83^ (v2.2).

Differential expression analyses were performed in each species independently and compared posteriorly, as illustrated in Figure S6. Bowtie2^84^ (v2.2.5), RSEM^85^ (v1.2.22) and DESeq2^86^ (p-value < 1e-3) were used to identify differentially expressed superTranscripts, using scripts from the Trinity package – just figure parameters were adapted. During analyses we identified a possible batch effect in samples from *T. angustula*: one nurse and one forager replicate were sequenced in different lanes and it seemed to affect sample correlation. This effect was corrected during differential expression analyses following the suggested protocol in DESeq2 documentation. No batch effect was identified in *B. terrestris* samples. To test whether any GO term was enriched in a set of differentially expressed superTranscripts compared to the total transcriptome, a classical Fisher’s exact test was performed using the R package TopGO^28^. Species comparisons of differentially expressed genes was based on gene annotation, using only unique and non-redundant terms (i.e. those genes not containing “uncharacterized protein” in their annotation). The list of overlapping genes was then manually curated to remove annotation incoherencies not detected computationally, i.e. when gene lists from both species were compared with our R script 18 terms were common, after manual curation we removed three genes from this list because of partial or redundant annotation matches (“transposase”, “transporter” and “cytochrome c oxidase subunit [fragment]”), leaving 15 genes in common. In the random sampling statistics this manual filtering correction was not used, so the numbers of common genes obtained with the computational comparison were used. Comparisons between the set of GO enriched terms and subgraphs was manual. The similarity network parameters was estimated with REVIGO^87^ using Medium (0.7) similarity threshold. In the interactive network mode of this program, the input data for Cytoscape^88^ was downloaded for further figure edition. Statistical tests of significance for comparisons were based on random sampling using R^89^ scripts, p-value smaller than 0.01 were considered significant. Scripts used are available at https://github.com/nat2bee/Foragers_vs_Nurses.

### DNA methylation analysis

Cleaning and adapter trimming of the bisulfite converted reads were performed using Trim Galore^90^ (v 0.4.3) wrapper script with default parameters. Complete transcriptome assemblies were used as reference so DNA methylation of coding regions could be analysed, since these regions are the main methylation targets in bees and other Hymenoptera^14^. PCR bias filtering, alignment of the cleaned reads and methylation call were performed using the BS-Seeker2^91^ (v 2.1.0), because this program allows the use of Bowtie2 in local alignment mode, which was necessary to properly align WBS reads to a transcriptome. CGmapTools^92^ (v 0.0.1) was used to filter low coverage methylated sites (< 10x) and to obtain DNA methylation statistics, including context use. Remaining statistical tests were performed using R, as follows: a random sampling test was used to verify whether the proportion of CG methylation found deviated from what was expected by chance; one-tailed z-test was used to test whether differences between the mean methylation observed in the set of superTranscripts was different from the general transcriptomic mean; and the correlation between methylation and gene expression was calculated using Spearman’s correlation coefficient between the superTranscript mean methylation and its normalized read count. Scripts used are available at https://github.com/nat2bee/Foragers_vs_Nurses.

## Acknowledgements

Authors would like to especially thank Dr. Yannick Wurm from Queen Mary University of London for his suggestions on this study and manuscript, they contributed immensely for this work. We also thank Dr. Isabel Alves-dos-Santos and Dr. Sheina Koffler from the Laboratório de Abelhas (University of Sao Paulo) for the support during *T. angustula* sampling, and Dr. Lars Chittka and Dr. Stephan Wolf from Bee Sensory and Behavioural Ecology Lab (Queen Mary University of London) for their support during *B. terrestris* sampling. Additionally, we would like to thank Dr. Bob Schmitz from the Schmitz lab (University of Georgia) for the support with DNA methylation sequencing, and to Susy Coelho from the University of Sao Paulo for technical assistance.

## Additional Information

This study was financed by FAPESP (São Paulo Research Foundation, processes numbers 2013/12530-4, 2012/18531-0 and 2014/04943-0), by CNPq (Conselho Nacional de Desenvolvimento Científico e Tecnológico, sponsorship to MCA), by Coordenação de Aperfeiçoamento de Pessoal de Nível Superior - Brasil (CAPES) [Finance Code 001], and developed at the Research Centre on Biodiversity and Computing (BioComp) of the University of Sao Paulo, supported by the university Provost’s Office for Research. Part of the bioinformatic analyses was performed at the cloud computing services of the University of Sao Paulo and of the Queen Mary University of London.

Authors declare they have no competing financial interests.

## Notes

https://github.com/nat2bee/Foragers_vs_Nurses

